# Quantifying heritability in functional regions of the coding and non-coding genome across diverse individuals

**DOI:** 10.64898/2026.10.30.763088

**Authors:** Theo V. Lupu, Harrison I.W. Wright, Camila Armirola-Ricaurte, Alicia Huerta-Chagoya, Andrew R. Wood, Josep M. Mercader, Alisa K. Manning, Micah R. Hysong, Inês Barroso, Caroline F. Wright, Michael N. Weedon, Robin N. Beaumont, V. Kartik Chundru, Gareth Hawkes

## Abstract

Most heritability studies have focused on common genetic variation and have largely excluded non-European or highly admixed populations. Here, we estimated the whole-genome heritability (h^2^_WGS_) of seven complex traits using 73,600,953 autosomal and chromosome X variants for 198,931 unrelated individuals from three genetic similarity groups. Adjusting for population stratification, we showed that rare variants (minor allele frequency<0.01) explain 30.6-67.7% of h^2^_WGS_. We used simulations to show how functionally-stratified heritability can be inflated due to linkage with missing or lower-quality variants. Finally, after accounting for confounding effects, we found substantial heritability enrichment from variants in non-coding RNA exons (maximum fold enrichment, FE=8.7, *P*=3.56e-7), UTRs (23.0, *P*=3.46e-30), distal regulatory regions (33.8, *P*=3.26e-42), and protein-coding exons (58.0, *P*=7.78e-15). Our study provides a framework for investigating the genetic architecture of complex human phenotypes using diverse whole-genome sequencing data and highlights the contribution of rare and non-coding genetic variants to the heritability of complex traits.

## Introduction

The genetic architecture of a phenotype describes the fraction of variance explained by genetic factors, which can be stratified across functional annotation categories, tissue types, and biological pathways^1–3^. Estimating the narrow-sense heritability of a trait using whole-genome sequencing (WGS) data (h^2^_WGS_) helps inform genetic architecture by quantifying the additive genetic variance in proportion to the phenotypic variance of a trait in a population, where additive genetic variance is increased by locus diversity and decreased by positive linkage disequilibrium (LD; describing the covariance between variants). Heritability is not fixed, and may vary in time, due to environment, and by genetic population. For example, a society where nutrition-rich food is depleted may present with a lower genetic heritability for standing height, i.e. where the environment limits genetic contributions to growth. There is also evidence for variation in heritability of body mass index (BMI) by age^4,5^ and for extreme BMI values^6^.

Due to population structure effects and variation in ancestral population allele frequencies^7,8^, commonly used heritability estimators also do not support the inclusion of non-homogenous populations, such that admixed individuals are routinely excluded from the analysis. Despite the increasing presence of admixed individuals in biobanks with genetic data, these populations remain empirically understudied in large genetic studies^9,10^, especially for h^2^_WGS_ estimates including rare and low-frequency variants (minor allele frequency (MAF)<0.01) or estimating the enrichment of different functional genomic categories^11–13^.

Existing studies have partitioned whole-genome heritability by broad functional annotation category (coding versus non-coding variants)^14,11^, or have focused on common variants using imputed data^14,11^ and summary-statistics-based methods such as LD score regression or SumHer^10–12^. Since h^2^_WGS_ considers overall genetic variation in a population sample at a given time, it tends to produce higher heritability estimates than the aggregated heritability from estimated effect sizes of lead variants in genome-wide association studies (GWAS; h^2^_WGS_) and includes variation due to low-LD variants not captured by imputation-based methods.

In this study, we aimed to answer three questions. First, how does genetic architecture and h^2^_WGS_ of complex polygenic traits differ between genetic similarity groups? Second, how do rare versus common variants contribute to heritability? Third, does the fold-enrichment of heritability in different functional variant categories, relative to the number of variants in each category, vary between complex traits? To answer these questions, we used 73,600,953 WGS single nucleotide variants (SNVs) and short indels from the All of Us research program in 198,931 genetically diverse individuals to investigate the genetic architecture of seven anthropometric and metabolic traits.

## Results

We stratified a total of 198,931 unrelated individuals with whole-genome sequencing (WGS) data from All of Us v8 (AoU) into genetically-inferred similarity groups if they had ≥60% probability of belonging to one group based on genetic similarity to super-population reference group individuals from the Human Genome Diversity Project^17^ and 1000 Genomes Phase 3^18^, as provided by AoU^19^ (**Methods; Supplementary Table 1**). Three groups had sufficient sample size for our analyses, individuals with genetic similarity to the African reference population (AFR; N=54,134), to the Americas reference population (AMR; N=59,831), and to the European reference population (EUR; N=84,966). We use the 3-letter reference population codes to refer to our own genetic similarity groups for brevity.

**Table 1:** Total h^2^_WGS_ meta-analysed across genetic similarity groups.

| **Trait** | **Total N** | **h<sup>2</sup><sub>WGS</sub>** | **95% CI** | **P** | **Heterogeneity P** |
|---|---|---|---|---|---|
| Height | 177,596 | 0.691 | 0.655, 0.728 | <1.00e-320 | 0.78 |
| BMI | 177,274 | 0.422 | 0.382, 0.461 | 3.06e-95 | 0.97 |
| WHR | 151,335 | 0.329 | 0.282, 0.376 | 7.69e-43 | 0.47 |
| WHRadjBMI | 149,623 | 0.306 | 0.252, 0.361 | 2.53e-28 | 0.31 |
| HGB | 83,929 | 0.343 | 0.137, 0.549 | 1.11e-3 | 1.14e-4 |
| T2D | 104,170 | 0.428 | 0.398, 0.459 | 2.00e-44 | 0.47 |
| HbA1c | 38,361 | 0.389 | 0.123, 0.654 | 4.11e-3 | 4.32e-2 |

*h<sup>2</sup><sub>WGS</sub>: narrow-sense whole-genome heritability, including autosomal and chromosome X variation; N: number of phenotypic records; WHR: Waist-hip ratio; BMI: body-mass index; WHRadjBMI: WHR adjusted for BMI; HGB: haemoglobin; T2D: type 2 diabetes mellitus; HbA1c: glycated haemoglobin.*
*Heterogeneity P<0.05: evidence for heterogenous total h<sup>2</sup><sub>WGS</sub> between genetic similarity groups.*

We considered 70,636,374 autosomal and 2,964,579 X-chromosome variants (SNVs and short indels) across genetic similarity groups. X-chromosome genetic relationship matrices (GRMs) have different variance-covariance structures to autosomal GRMs and were thus treated as a separate, but jointly-modelled, contribution to heritability (**Supplementary Note 1**)^20,21^. To minimise error due to population structure from recent admixture, we classified each variant category by MAF (rare: 1e-4≤MAF<0.01 versus common: 0.01≤MAF<0.5) and per-variant LD score, calculated within genetic similarity groups. As variant frequencies were estimated separately for each genetic similarity group, we analysed different rare and common variants per group, and different sets of rare and common variants were shared between groups (**Supplementary Figure 1**). These patterns reflect recent admixture patterns and the larger genetic diversity in non-European populations. We additionally included 120 genetic group-specific common-variant PCs as covariates alongside age, age squared, sample source (whole blood or saliva), and genetic sex (**Methods**).

### Total and allele frequency-partitioned h^2^_WGS_ is consistent across genetic similarity groups

For each genetic similarity group, we quantified the total heritability (h^2^_WGS_) and the proportion of heritability attributed to variants with rare or common MAFs (h^2^_R_ and h^2^_C_ respectively). We used MINQUE for Partitioning Heritability (MPH)^22^ in seven well-measured anthropometric and metabolic traits height, body mass index (BMI), waist-to-hip ratio (WHR), WHR adjusted for BMI (WHRadjBMI), haemoglobin (HGB), glycated haemoglobin (HbA1c), and type two diabetes mellitus (T2D).

We found that meta-analysed h^2^_WGS_ (**Table 1**; **Figure 1A**) was not significantly different across genetic similarity groups for anthropometric traits and T2D (mean heterogeneity *P=*0.403 across all traits; **Supplementary Table 2**). However, we observed significant differences between EUR and AMR/AFR h^2^_WGS_ for blood-cell traits: HGB (HGB: AFR h^2^_WGS_=0.304, AMR h^2^_WGS_=0.529, EUR h^2^_WGS_=0.192, heterogeneity *P=*1.14e-4), and between AFR and EUR/AMR for HbA1c (HbA1c: AFR h^2^_WGS_=0.117, AMR h^2^_WGS_=0.595, EUR h^2^_WGS_=0.422, heterogeneity *P=*4.32e-2).

**Figure 1:**
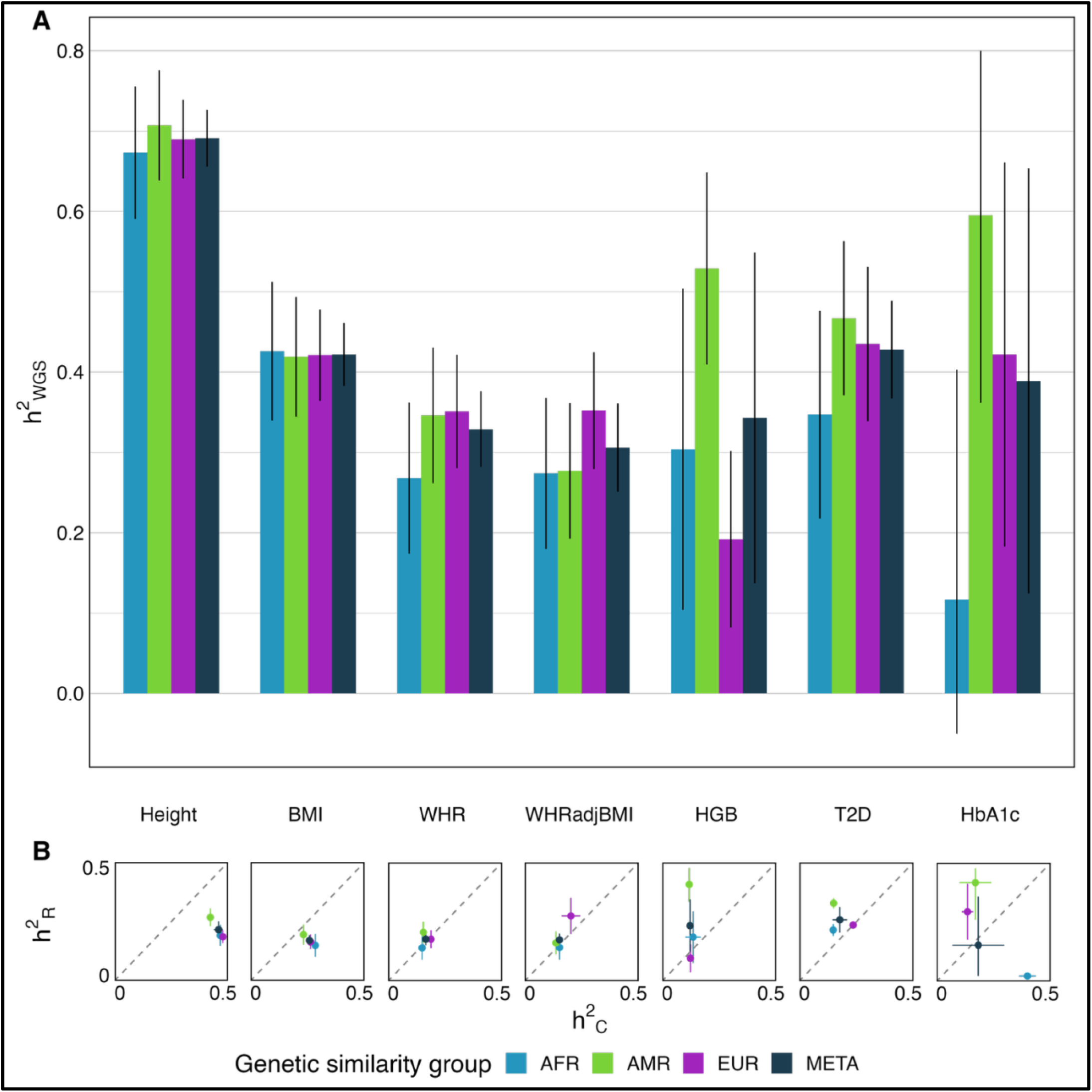
Heritability of seven traits across genetic similarity groups. **(A)** Total heritability (h^2^_WGS_) per trait, including autosomal and chromosome X variation. **(B)** Rare (1e-4≤MAF<0.01; h^2^_R_) versus common (0.01≤MAF<0.5; h^2^_C_) heritability per trait, including autosomal and chromosome X variation. Dotted line is y=x and error bars represent 95% confidence intervals (±SE*1.96), which are truncated at axis limits (0≤h^2^≤0.5).

The percent of meta-analysed rare heritability out of total estimated heritability (h^2^_R_/h^2^_WGS_) ranged from 30.6% (height) to 67.7% (HGB). This ratio was similar between genetic similarity groups for BMI (heterogeneity *P=*0.351), WHR (*P*=0.567), and WHRadjBMI (*P=*0.874), but was significantly heterogenous for height (*P=*5.79e-3), HGB (*P*=2.53e-3), and T2D (*P*=7.78e-40) (**Figure 1B**; **Supplementary Figures 2&3**). The strongest evidence for heterogeneity between genetic similarity groups was attributable to the magnitude of contribution of rare variants to T2D heritability (h^2^_R_/h^2^_WGS_=61.0% (AFR), 72.0% (AMR), 54.0% (EUR); *P*<9.06e-39). Despite HbA1c showing heterogeneity between genetic similarity groups for total heritability, the ratio of rare to common h^2^ was not significantly heterogeneous (*P*=0.052), likely due to REML model mis-specifications regarding rare-variant allelic effect distributions in the AFR cohort (**Discussion**).

Variants on chromosome X have been previously reported to have large effects on HbA1c, and thus T2D diagnosis^23,24^ (e.g. *G6PD* in AFR-like individuals). We found that common variants on chromosome X contributed most significantly to HbA1c and T2D heritability for the AFR group (X-chromosome common variant HbA1c h^2^_C-X_=0.131, *P*=1.09e-20; T2D h^2^_C-X_=0.016, *P=*4.66e-3). Including chromosome X variation corresponded to a total h^2^_C_ increase of 31.7% for HbA1c and 11.4% for T2D in AFR, compared to h^2^_C_ attributable to autosomal variation alone. The contributions of chromosome X variants to h^2^_C_ in AMR and EUR for HbA1c and T2D were more modest (an increase in h^2^_C_ ranging from 0.36% to 7.22%; **Supplementary Table 2**). In comparison, the contribution of rare autosomal variants to AFR HbA1c h^2^_WGS_ estimates were significantly negative (h^2^_R-auto_=-0.234, *P* = 9.82e-4), the only example of a significantly negative h^2^ component in our analysis, resulting in a total heritability which was not statistically significantly different from zero (h^2^_WGS_=0.117, *P=*0.12). Negative heritability estimates from MPH indicate low signal-to-noise ratios or model misspecifications, such as the presence of non-additive genetic effects or strong genotype-environment interactions. Chromosome X variants also had a smaller contribution to heritability for other traits and genetic similarity groups (**Supplementary Table 2**).

### The heritability of complex traits is enriched in noncoding RNA exons, protein-coding exons, UTRs, and distal regulatory regions

We next partitioned rare and common variants by functional annotation to test whether the contribution of functional variant categories to h^2^_WGS_ were enriched relative to the number of variants contained therein, and whether this enrichment differed between traits and genetic similarity groups. We did not include chromosome X in our functionally-stratified analyses as X-GRMs cannot be combined with autosomal GRMs, and functionally stratified X-GRMs would be too noisy given the small number of variants which would be used to construct them **(Supplementary Note 1)**. In the following sections, total h^2^_WGS_ is therefore estimated from autosomal variants only.

We split autosomal variants into seven functional categories using Ensembl’s Variant Effect Predictor^25^ (**Methods**; **Supplementary Table 3**): protein-coding, non-coding RNA exons (ncRNA), untranslated regions (5’UTRs plus 3’UTRs) and gene-proximal variants (±5kb around the UTRs), introns (including protein-coding gene and ncRNA introns), predicted intergenic regulatory regions, and variants not otherwise annotated. Any variant which was annotated to more than one category was assigned to its first allocation in the given list.

Across traits, the largest proportion of heritability (mean=47%, SD=22%) was attributed to intronic regions (**Figure 2A**; **Supplementary Tables 4&5**). However, variant annotation category sizes were not equal (**Supplementary Table 3**), with intronic and un-annotated categories containing the highest number of variants (29.43% and 26.28% of autosomal variants, respectively, averaged and weighted by genetic similarity group size). Similarly, 79% of autosomal variants considered were rare, but contributed on average only half the total heritability (50%; SD=12%). Therefore, to quantify the contribution of each functional annotation category to genetic heritability, relative to the number of variants contained therein, we calculated per-category fold enrichment (FE; **Methods**). We define significant enrichment as FE>1 (P<0.05), and significant depletion as FE<1 (P<0.05).

**Figure 2:**
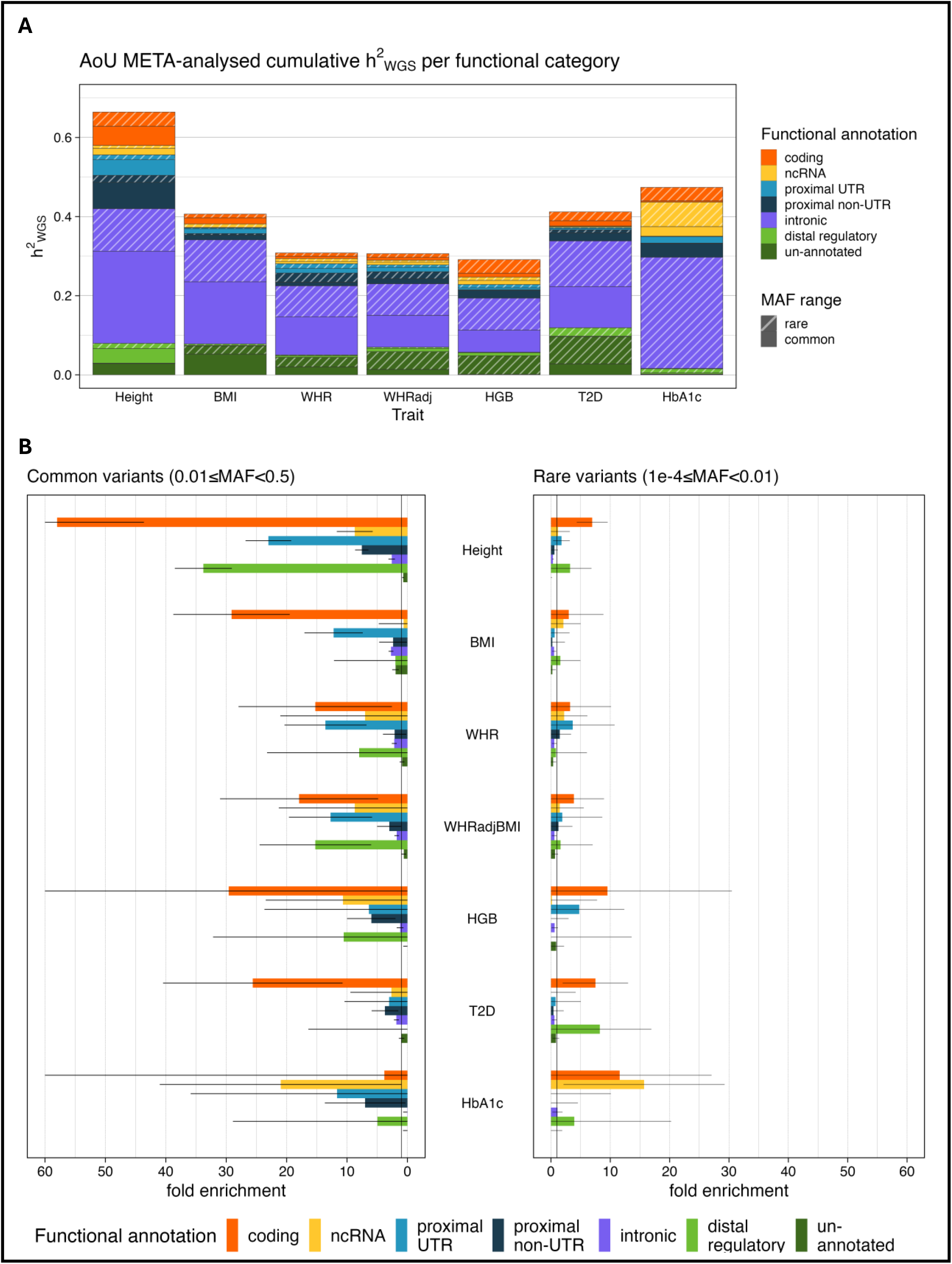
Functionally partitioned meta-analysed heritability of seven traits. **(A)** Meta-analysed cumulative autosomal h^2^_WGS_ per functional category. Shaded bars represent rare-variant (1e-4≤MAF<0.01) heritability, unshaded bars represent common-variant (0.01≤MAF<0.5) heritability. **(B)** Meta-analysed fold enrichments per functional annotation and MAF category, per trait. Fold enrichment (FE) = (h^2^_Cat_ / ∑h^2^_WGS_) / (m_cat_ / ∑m), where h^2^_Cat_ / ∑h^2^_WGS_ = proportion of heritability and m_cat_ / ∑m = proportion of variants per category. Meta-analysed FE corresponding to heritability<0 are not presented. Error bars represent 95% confidence intervals (FE±1.96*SE) and are truncated at axis limits (0 ≤FE ≤ 60).

Functional annotation groups with statistically significant contributions to h^2^_WGS_ tended to also have statistically significant FE values (based on -log_10_(*P*) Spearman’s correlation, ρ=0.687, *P*<2.2e-16). Conversely, h^2^_WGS_ magnitude was more modestly correlated with FE magnitude (ρ=0.237, *P*=0.019), since FE is a function of both heritability and annotation category size (**Supplementary Figure 4**). This pattern follows when considering h^2^ due to rare and common variants separately (h^2^_R_: log_10_(*P*) ρ=0.574, *P*=2.29e-5; h^2^_WGS_ versus FE ρ=0.309, *P*=0.031; h^2^_C_: log_10_(*P*) ρ=0.746, *P*=1.16e-9; h^2^_WGS_ versus FE ρ=0.133, *P*=0.360).

We found the highest overall enriched contribution to heritability in protein-coding exons, ncRNA exons, UTRs, and predicted distal regulatory regions. Common variants on average presented with a larger FE than rare variants per functional annotation bin (**Figure 2B**; **Supplementary Tables 6&7**). More specifically, in exonic regions we found enrichment due to common and rare ncRNA variants (common-variant ncRNA FE=8.72 for height and rare-variant ncRNA FE=15.7 for HbA1c) and across traits due to protein-coding variants (ranging from to FE=6.96 to 45.3). Common proximal UTR variants were also enriched for anthropometric traits (FE=12.2 to 23.0), while common proximal non-UTR variants were enriched for height, HGB, and HbA1c (ranging from FE=3.72 to 7.56).

We found modest enrichment in common intronic regions for anthropometric traits (ranging from FE=1.78 to 2.72), with significant depletion in rare intronic regions for height, BMI, WHR, and T2D (ranging from FE=0.33 to 0.59). Distal predicted regulatory regions were enriched for height (FE=33.8) and WHRadjBMI (FE=15.2). Finally, we detected significant fold enrichment due to common un-annotated variants only for BMI (FE=1.96) with depletion for height, BMI (considering common variants), and WHR (ranging from FE=-0.023 to 0.71). Of these, there was significant heritability (h^2^>0, P<0.05) only due to common un-annotated variants for height. There were few cases of per-category heterogenous FE (*P*<0.05) across genetic similarity groups (**Supplementary Table 6**), most significantly common-variant intronic FE for height (P= 5.58e-4).

### Functional category sub-divisions of protein-coding and noncoding RNA exonic variants reveal additional enrichment patterns

To determine which variant function subtypes drove the enriched contribution of exonic variants (protein-coding and ncRNA) to heritability, we sub-divided those categories (**Figure 4**; **Supplementary Table 3**). We split protein-coding variants into predicted loss-of-function (pLoF; 2.51% of coding), splice-region (7.69%), missense (53.07%), and synonymous (36.59%) variants, and split ncRNA into long noncoding RNA (lncRNA; 98.49% of ncRNA) and other ncRNA including miRNA (0.03%), rRNA (0.17%), snoRNA (0.35%), and snRNA (0.64%) variants. To reduce the probability of falsely assigning enrichment to a functional category only due to our prior assumptions of importance, and because each sub-category was increasingly small, we additionally allowed variants to be assigned to multiple categories (**Methods**). The overlap between protein-coding and ncRNA exons was minimal: an average of 1.64% of variants in any ncRNA sub-category overlapped with any protein-coding sub-category. To account for the overlap, we adjusted h^2^_WGS_ and enrichment results reflecting the per-variant contribution to GRMs (**Methods**).

We found significant enrichment for the contribution of protein-coding sub-categories to h^2^_WGS_ (**Figures 2B&3A**; **Supplementary Tables 7&8**). Rare and common missense variants were enriched for height, BMI, HGB, and T2D (ranging from FE=2.13 to FE=21.7, *P*<0.05), as were common splicing-altering variants for height (FE=46.6, P=3.19e-3). The largest fold-enrichment was for common-variant pLoFs for HbA1c (FE=638.1, *P*=1.76e-2), however, there were fewer than 2,000 pLoF variants with MAF>0.01 per genetic similarity group, making for potentially noisy GRMs and overestimated enrichment values. We also observed evidence of heterogeneity between genetic similarity groups (*P*<0.05) in FE attributable to coding variants, specifically missense variants, for height.

Across genetic similarity groups, heritability due to ncRNA exons was significantly enriched (**Figures 2B&3B**; **Supplementary Tables 7&9**) for height (FE=8.72, *P*=3.56e-7) and HbA1c (FE=15.7, *P*=3.39e-2). Specifically, common-variant lncRNA variants were enriched (FE=1.51, *P*=9.41e-7 for height; FE=10.2, P=4.60e-2 for HbA1c).

We hypothesised that the observed enrichment in synonymous-variant heritability could be due to linkage with missing or poorly genotyped causal protein-altering variants. To test this, we performed simulations for the AFR cohort (**Figures 3C&D**; **Supplementary Tables 10&11**; **Methods**). We chose to simulate only using the AFR cohort as it is the smallest group of three thus minimising computing resources required, whilst ensuring the results would be robust to increased population structure relative to EUR-like cohorts. Briefly, per MAF-LD bin, we sampled Normally-distributed variant effect sizes scaled by MAF such that the simulated per-bin h^2^_WGS_ replicates the per-bin h^2^_WGS_ ratios observed for the real phenotypes. From these effect sizes, we generated a simulated phenotype with added random noise to match the observed total h^2^_WGS_. We used the GRMs from our observed data.

**Figure 3:**
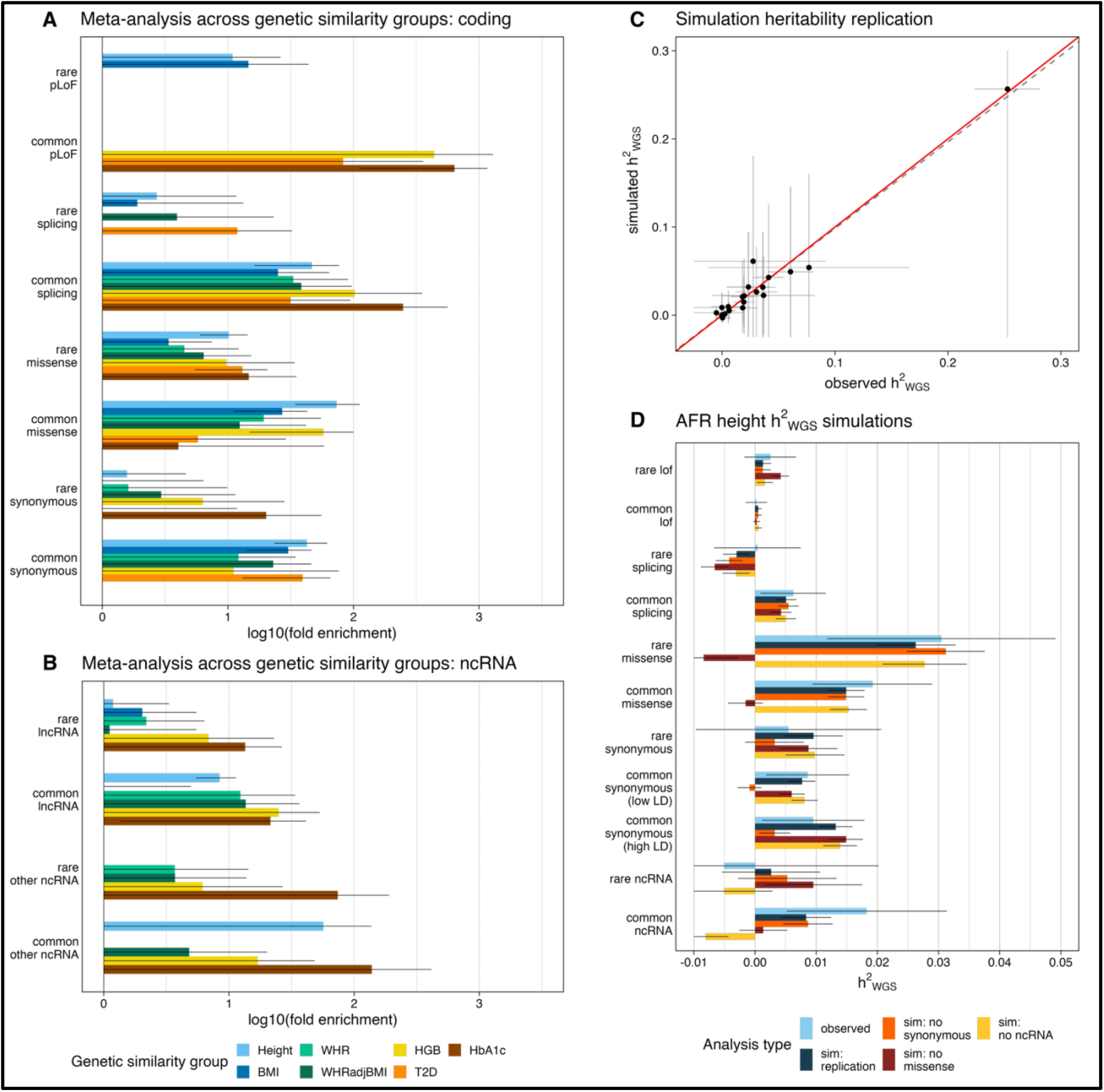
Fold enrichment in functional annotation sub-categories. Fold enrichment due to coding **(A)** and ncRNA **(B)** sub-categories, meta-analysed across genetic similarity groups. Fold enrichment (FE) = (h^2^_Cat_ / ∑h^2^_WGS_) / (m_cat_ / ∑m), where h^2^_Cat_ / ∑h^2^_WGS_ = proportion of heritability and m_cat_ / ∑m = proportion of variants per category. Dotted grey line is the enrichment threshold (FE=1). **(C)** Simulations replicating empirical AFR height h^2^_WGS_ results across functional categories. Red line is y=x, dotted grey line is line of best fit (adjusted R-squared=0.92, P=2e-24); **(D)** Simulated h^2^_WGS_ in coding and ncRNA categories, comparing observed AFR height h^2^_WGS_ (blue) with simulated AFR phenotypes replicating h^2^_WGS_ per bin (dark blue), and simulated AFR phenotypes replicating h^2^_WGS_ per bin while equating heritability in either synonymous (orange), missense (brown), or ncRNA (yellow) bins to 0, while keeping total h^2^_WGS_=0.673. Simulation estimates per analysis type are meta-analysed across 10 repeats. For **(A), (B)**, and **(D)**, error bars represent log-transformed 95% confidence intervals (log10[FE ± 1.96*S])), and for **(C)** error bars represent ±1.96*SE. Error bars are truncated at axis limits (log10[1] ≤ FE ≤ log10[3000]).

Our simulations accurately replicated the observed h^2^_WGS_, meta-analysed across 10 repeats (adjusted R-squared=0.92, *P*=2e-24; **Figure 3C**). We then set h^2^ from synonymous variants to 0 in our simulations (**Figure 3D**) while keeping total heritability the same (h^2^_WGS_=0.673). This reduced simulated common-variant synonymous h^2^ and fold enrichment from h^2^=0.021 (*P*=3.39e-34) and FE=45.0 (*P*=5.28e-32) to h^2^=0.002 (*P*=0.148) and FE=5.20 (*P*=0.152) respectively. Specifically, common-variant high-LD synonymous h^2^ and fold enrichment decreased from h^2^=0.013 (*P*=1.89e-22) and FE=56.2 (*P*=3.23e-21) to h^2^=0.003 (*P*=1.30e-2) and FE=13.8 (*P=*1.57e-2). Comparing these FE values for the replication simulation set versus when synonymous heritability=0 indicates that approximately one quarter (24.6%) of the simulated enrichment could be due to haplotype effects. In contrast, setting h^2^ from other significantly enriched functional categories to 0 resulted in null heritability and no enrichment (**Supplementary Tables 10&11**; **Supplementary Figure 5**). These data suggest that only the observed synonymous h^2^_WGS_ contribution and enrichment signals are likely to be inflated by haplotype effects.

## Discussion

In this study we used >70M whole-genome sequenced variants from 198,931 diverse individuals to investigate the contribution of rare and common variants to the heritability of seven anthropometric and metabolic traits across genetic-similarity groups. We also quantified as fold-enrichment (FE), the relative contribution of different functional genomic categories to heritability. Firstly, we found that rare variants significantly contribute to heritability, constituting 30.6-67.7% of total heritability, but that this proportion varies substantially by trait. Second, we found that fold enrichment (FE) for the contribution of rare and common variants to heritability was similar across most functional annotation categories and genetic similarity groups, but rare variants were less enriched than common variants. We report a substantial enrichment for the contribution of non-coding annotations to heritability estimates, particularly noncoding RNA (ncRNA), untranslated regions (UTRs), and distal regulatory regions.

Our results showed largely homogeneous total h^2^_WGS_ across genetic similarity groups, with some differences observed for HbA1c, HGB, and T2D. In the broader population context, these differences could be due to biased h^2^_R_ in highly admixed groups such as AMR. Additionally, if environmental factors explain more of the trait variance, genetic contributions will proportionately decrease, and changing environment may result in changing heritability estimates over time. Given that most of the traits we considered were ascertained in electronic health records, there could also be socioeconomic or geographical biases in which individuals have access to different health resources and care. Correspondingly, large variations in non-genetic factors affecting a phenotype may cause large signal-to-noise ratios, which can either underestimate heritability or result in large uncertainty.

Additionally, rare variants which affect trait heritability may be found at different allele frequencies between populations. This can affect both the proportion of rare versus common heritability between genetic similarity groups, and total heritability estimates in cohorts where there are low numbers of carriers of the causal variants. There are examples of AMR-specific variants with large effects on T2D and HbA1c measures^26^, as well as adaptive phenotypes enriched in AFR-like genetic similarity groups and subsequently admixed populations that affect red blood cell shape or lifespan due to their protective effects on malarial infection^10,27–29^, including sickle cell anaemia, G6PD deficiency, haemoglobin C, or thalassemia. One such example is the X-chromosome *G6PD* missense variant *rs1050828* (with minor allele frequencies (MAF) of 13% in AFR, 1% in AMR, and 0.01% in EUR), which is associated with lowering HbA1c levels independently from glucose^30–32^, and consequently with a risk for T2D underdiagnosis^24^. We found a significant h^2^_WGS_ component attributed to common X-chromosome variants only in AFR for HbA1c and T2D, with low X-chromosome contributions for other traits or genetic similarity groups. Fully capturing the contribution of *rs1050828* or other large-effect but common variants to h^2^_WGS_ would require modelling them separately as fixed effects to preserve the assumption of effect size Normality for variants included in GRMs^33^.

We observed some evidence of statistical instability of model-misspecification in our results, exemplified by a statistically significant negative rare-variant heritability contribution to HbA1c in AFR individuals. Wainschtein et al.^34^ found that large GRM diagonal values (reflecting population structure and haplotype effects, sequencing artefacts, or ultra-rare variants carried by an individual) can downwardly bias heritability estimates. A similar result was reported by Liu et al.^35^ when investigating rare pLoF variants with non-Gaussian effect sizes distributions. The potential biases outlined are likely maximised by the small number of phenotype records for both HbA1c and HGB.

Small ncRNAs such as *RNU4-2* are known to affect rare developmental disorders^36,37^, but our work highlights the importance of ncRNA variants for common phenotypes. We detected significant enrichment in the contribution to h^2^_WGS_ from common ncRNA variants for height and HbA1c which was attributable to lncRNA variants. Supporting this, there are several well-known distal regulatory and ncRNA variants involved in imprinting control regions for anthropometric traits^38,39^ such as *IGF2, CDKN1C*, and the lncRNA growth suppressor *H19*. Some lncRNAs are also known to affect HbA1c directly^40,41^, and others are associated with T2D via differential expression in pancreatic islets, beta cell programming^42^, insulin resistance^43^, or adipose tissue distribution^44^. However, we did not observe an enrichment for the contribution of ncRNA to T2D, which could be due to reduced sensitivity when considering a binary trait on the liability scale.

Our data also shows significant enrichment of heritability in proximal and distal regulatory variant categories. Enrichment in UTR regions was similar across traits, while enrichment for distal regulatory variant heritability was high for height and WHRadjBMI for common variants. Previous research has identified UTR variants affecting several traits^45^ and rare aggregate associations in non-coding gene-proximal UTRs have been linked to height^46^. Our results also suggest that the fraction of heritability of anthropometric traits attributed to distal regulatory elements is highly concentrated in common variants. Supporting this, previous common-variant height GWAS have found large enrichment in regulatory variants^47,48^, specifically functional chromatin annotations.

Finally, un-annotated and intronic variants form a large part of heritability for most traits but are only moderately enriched across several traits. This small enrichment despite high h^2^_WGS_ could suggest that our annotation method could be imperfect. Alternatively, un-annotated and intronic variants may have small individual effects and contribute significantly to heritability only in aggregate as they form a large part of the genome.

We also found significant fold-enrichment contribution to heritability for protein-coding variants down to MAF>1e-4 across traits in missense variants and those with known splicing effects. As missense and splice-altering variants affect protein sequence, it is likely that many have some effect on function. On the other hand, most known and predicted LoF variants are ultra-rare (below our threshold of MAF<1e-4) and fewer in number^49,50^ compared to variants with other coding consequences. Therefore, GRMs built with pLoF variants are likely to be very noisy (**Methods**) and non-representative of true pLoF genetic variation. We believe that the inability to capture the contribution of these true causal ultra-rare pLoF variants is why our calculated pLoF enrichment signals are smaller than for other coding sub-categories.

A result that likely does not reflect true genetic architecture is the enriched contribution of common synonymous variants to heritability. Previous studies have found no conclusive evidence supporting true synonymous variant effects on trait heritability in any organism (reviewed by Oelschlaeger^51^), with some cited potential small effects on alternative splicing, transcription regulation, or mRNA stability. We note that REML heritability enrichment in a given GRM does not directly indicate genotype-phenotype causality, as REML uses maximum likelihood to assign heritability to the variance component that most stably associates with variation in a phenotypic trait^20,52^. This method favours GRMs created with variants under little or no selection pressure, like synonymous variants, as these variants are numerous and with higher MAF so would generate less noisy GRMs than the other coding sub-categories like pLoFs. Given their proximity, however, synonymous variants are also likely in high-LD with true causal protein-coding variants. Our simulations indicated that at least 24.6% of the observed synonymous enrichment is likely due to haplotype sharing with such protein-altering variants.

There are some limitations to our study. Firstly, although we took conservative approaches to correct for population stratification, with strict sample filtering, including 120 genetic principal components, and using MPH which is designed to accommodate cohorts with larger haplotype blocks, there may still be fine-scale population structure effects which will introduce some bias into the estimates. Another source of potential bias is assortative mating, which was not accounted for in our analysis, and would be expected to increase estimates of heritability. Finally, REML-based heritability alone serves as an indicator of where causal variants may lie but does not directly explain genotype-phenotype causality. Difficult to sequence regions, including regions susceptible to segmental duplications, low complexity regions, and tandem repeat regions, will not be captured by short-read sequencing technologies. The effects of causal variants in these regions will however be captured in part through LD with variants that are sequenced, and these captured variants may not be in the same assigned functional category, resulting in misleading enrichments in some categories. We discuss these issues further in **Supplementary Note 2**.

In conclusion, our results support further analysis to quantify the contribution of functional variant categories to genetic heritability, the testing of rare variants in aggregate for association studies, and the inclusion of individuals of diverse and admixed populations in heritability and association analyses.

## Acknowledgements

TVL, HW, RNB, and MNW were supported by Medical Research Council (MRC) grant MR/Y003748/1. GH was supported by the MRC grant UKRI327. VKC and CFW were supported by the Wellcome Trust grant 226083/Z/22/Z. ARW was supported by the Academy of Medical Sciences, the Wellcome Trust, the Government Department of Business, Energy and Industrial Strategy, the British Heart Foundation and the Diabetes UK Springboard Award (no. SBF006\1134). This study was supported by the National Institute for Health and Care Research Exeter Biomedical Research Centre. The views expressed are those of the authors and not necessarily those of the NIHR or the Department of Health and Social Care. We gratefully acknowledge *All of Us* participants for their contributions, without whom this research would not have been possible. We also thank the National Institutes of Health’s *All of Us* Research Program for making available the participant data examined in this study.

## Data Availability

Individual-level data cannot be shared publicly because of data availability and data return policies of the *All of Us* Research Platform. Data are available from *All of Us* for researchers who meet the criteria for access (https://allofus.nih.gov/).

## Code Availability

The code used to run heritability analysis, as well as the code used to generate figures, is available at [TO BE ADDED UPON ACCEPTANCE].

## Conflicts of Interest

The authors have no conflicts to declare.

## Methods

### WGS datasets

The whole genome sequencing performed for *All of Us* v8 (AoU) had average coverage 37.9X using Illumina NovaSeq 6000 sequencing technology. The genome build used for mapping was GRCh38: single variant nucleotide variants and short indels were jointly called using DRAGEN 3.7.8. The full AoUv8 WGS data was provided exclusively in HAIL VDS format, with allele-level VETS filtering flags ^53^, which we filtered on. We exported the Hail VDS to VCF format using Hail v.0.2.134-952ae203dbbe. We applied bcftools^54^ to these vcfs to split multi-allelic sites, and left-aligned all REF and ALT alleles using a 1000G b38 reference available at https://ftp.1000genomes.ebi.ac.uk/vol1/ftp/technical/reference/GRCh38_reference_genome/ (accessed 30/03/2024). We assigned all variants a unique ID (CHR:POS:REF:ALT), followed by the application of plink2^55^ to convert each filtered pVCF shard to plink2 format, and finally concatenated those plink-shards into per-chromosome p(gen/sam/var) files.

### Defining sets of unrelated individuals

We used genetic similarity groups defined by the *All of Us* Research Program (see AoUv8 QC report Appendix G^19^), which were stratified into groups based on genetic similarity to harmonised continental metadata from the Human Genome Diversity Project^17^ and 1000 Genomes Project training data^18^. We dropped individuals with known quality issues and removed related individuals to limit shared environmental effects using KING coefficients^56^, keeping KING≤0.0221=less than fourth degree relatedness. The maximum independent set of unrelated individuals were selected per genetic similarity group using the `maximum_independent_set()`function from the NetworkX Python package^57^. After QC and kinship filtering, we retained the following samples: individuals with >60% genetic similarity to the African American/Black Caribbean-like reference panels (AFR; N=54,134), to the admixed American-like group (AMR; N=59,831), and to the European-like group (EUR; N=84,967) (**Supplementary Table 1**).

### Phenotypes and Covariates

Individual-level phenotypes and covariates were obtained from AoU (see **Supplementary Table 1** for OMOP concept IDs). T2D controls were defined as individuals with no previous diagnosis or self-reporting of any diabetes or related conditions (prediabetes, gestational diabetes, or glycosuria), had no prescription of glucose-lowering diabetes medication, and had maximum HbA1c<5.7%, maximum random glucose<200 mg/dL, and maximum fasting glucose <100 mg/dL. T2D controls and cases were further filtered for age>45. For each genetic similarity group, we rank-inversed all continuous phenotypes per sex, re-combined the results, and residualised on all covariates (120PCs, age and age squared, sample source, and sex).

### Per-variant linkage disequilibrium

We computed whole-genome per-variant linkage disequilibrium (LD) scores per final genetic similarity groups using the GCTA Software tool v.1.94.1^58^. The per-variant LD statistic is the sum of squared correlation between that variant *m* and neighbouring variants *k* within a 1Mb window with MAC>5, excluding pairs with r^2^ >0.01:

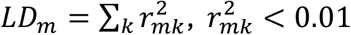

We included rare variants in our LD calculation to capture trends in population stratification and recent admixture. To maximise computational efficiency for this step, we divided the genome into 20Mb consecutive blocks with 1Mb overlaps (AFR, AMR) or 2Mb blocks with 500KB overlaps (EUR, EUR), depending on sample size. Per variant, we retained the maximum LD score calculated using this overlap method.

### GRMs and heritability estimation assumptions

We used MPH v.0.55.0^22^ to estimate genetic heritability per genetic similarity group. This method estimates the proportion of phenotypic variance explained by genetic variance and covariance between individuals described in several GRMs. MPH uses an iterative MINQUE process which mirrors Fisher-scoring GREML (Genome-based Restricted Maximum Likelihood) used by the GCTA Software tool^58^ with additional optimisation to escape common GREML convergence failures. It additionally uses stochastic trace estimation and parallel computing to decrease computational time and memory usage. MPH was originally developed for animal breeding h^2^_WGS_ estimation, on the premise that modern fast approximation methods for estimating variance components (e.g. HE and LD-score regression) cannot effectively account for related individuals or complex LD structures.

To account for potential in-sample dependency of per-variant effect sizes on LD score^15,59^ we performed GREML-LDMS (MAF and LD Stratified GREML) with multiple MAF and LD stratified GRMs (**Supplementary Table 3**) and the `--reml-no-constraint`flag similarly to GREML-LDMS-I in Evans et al.^52^ and Wainschtein et al.^34^. GRMs were calculated with GCTA (`--make-grm-part`, `--make-grm-alg 0`) and converted to the MPH format by incorporating the number of variants used to calculate each pairwise relationship into the GRM grm.bin file.

The standard Van Raden 2 model used by GCTA and MPH for h^2^_WGS_ estimation assumes an infinitesimal model of Normally distributed independent random effects, where each variant contributes additively to the total h^2^_WGS_. Generally,

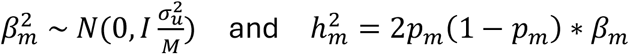

where *β*_*m*_ is the effect size of variant *m*, 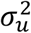 is the variance of per-variant effects, 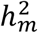 is the per-variant heritability, *p*_*m*_ is the allele frequency, and *M* is the total number of Variants per bin. Following these assumptions, h^2^_WGS_ is equal to total genetic heritability if all causal variants are included in the analysis^20,22^.

Thus, the GRM value describing genetic covariance between individuals *i, j* for *M* common variants *m* is:

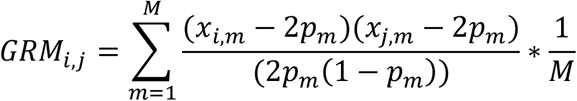

where the numerator centres the individuals’ genotype values and the denominator normalises allele frequencies. Off-diagonal values correspond to the relationship between individuals, expected < 1, while diagonal values normally ∼1. However, such GRMs computed using mainly rare variants have a diagonal values >>1, since 1/2*p*_*m*_(1 − 2*p*_*m*_) grows exponentially with MAF. This means that rare-variant GRMs can be imprecise due to the sequencing errors which occur more often for rare variants. Sequencing depth or total number of rare allele differences between individuals can also cause significant variation in diagonal values in the GRM, where individuals with larger corresponding diagonal values contribute more to heritability. Additionally, it has been shown that low-MAF Variants can have a upwardly biased effect size^60,61,33^ in h^2^_WGS_ analysis by over-accounting for negative selective pressure. To limit these biases, we modified GCTA to scale per-variant contributions to the GRM akin to the LDAK model. We exponentially downweighed per-variant contribution for lower-MAF Variants:

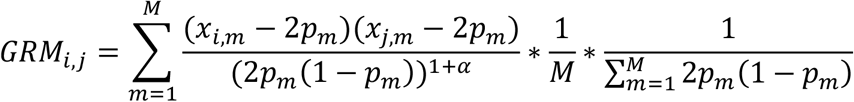

We used *α* =-0.25 as recommended by Speed et al.^33^. To reflect the addition of alpha, we normalised by 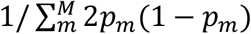 additionally to 1/*M*.

### Defining variant bins by functional annotations, MAF, and LD

We aimed to estimate WGS heritability (h^2^_WGS_) and enrichments by functional annotation, so we divided autosomal Variants by non-overlapping functional category using Ensembl’s Variant Effect Predictor v.115.1^25^. To create GRMs per genetic similarity group, we divided each functional category by MAF and then LD **(Supplementary Table 3**). We stratified the genome into seven functional categories based on prior studies investigating summary statistics-based partitioned heritability in White British Europeans^33,45,62^. To avoid category overlaps, variant lists were filtered using the following hierarchy: protein-coding exons > ncRNA exons > proximal (UTR) > proximal (non-UTR) > introns > distal constrained regulatory > un-annotated. Annotation MAF bins were rare (1e-4<MAF≤0.01) and common (0.01<MAF≤0.5). LD bins were divided into two groups using the median variant LD r^2^ score per MAF bin of each annotation. We also created four separate GRMs for the X-chromosome per genetic similarity group (same MAF and LD divisions as above) to include when calculating the heritability of traits that have significant GWAS X-chromosome variance components, such as HbA1c measurements for individuals of African descent with the *G6PD* rs1050828 missense variant^23,30^.

We further tested the partitioned heritability by separating the protein-coding exon annotation (per consequence: predicted loss of function, splicing, in-frame missense, synonymous, and other) and the ncRNA exon annotation (per ncRNA type: lncRNA and other ncRNA, including miRNA, rRNA, snoRNA, and snRNA). From our list of synonymous variants, we specifically removed all non-MANE transcript variants annotated as transcription factor binding sites by VEP, with predicted high splice donor or acceptor loss or gain likelihood (SpliceAI^63^ score≥0.2, adding them to the splicing variant list) scores or high predicted deleteriousness (combined annotation-dependent deletion score, CADD^64^ score≥25, adding them to the ‘other coding’ variant list) scores. Enrichments for ncRNA sub-categories were adjusted to account for overlaps and per-variant weights by taking the cross-product of the variant incidence matrix (noting which variants belong to which category) with the SNP weighing matrix (reflecting per-variant contribution to GRMs) to apply the MPH function recalculate_enrichments(). When calculating partitioned heritability for these sub-categories, we aggregated all non-coding Variants, non-RNA into one functional annotation category to retain power.

When aggregating MPH heritability across *j* LD and/or MAF bins *i*, we accounted for potential covariance between groups by recalculating standard errors *SE*_*j*_ as:

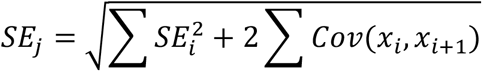

Where *SE*_*j*_ =total standard error for h^2^_WGS_ estimation in bin *j*, ∑ *SE*^2^= summed variances for all *i* bins in *j, Cov*(*x*_*i*_, *x*_*j*_)=covariance between MAF-LD bins. We did not apply this to total genome-wide narrow-sense heritability since all methods provided a total h^2^_WGS_ score. To maximise power, we also conducted meta-analyses for partitioned trait heritabilities and enrichments across genetic similarity groups using the random effect model with inverse variance weighing in the R package metafor^65^. To examine correlations between variant functional categories, we normalised trait heritability per functional category across meta-analysed results so that total heritability=1.

### Fold enrichment

We recalculated fold enrichment and standard errors (SE) following the MPH source script after combining results from the two LD bins per MAF-annotation category. We recalculated fold enrichment per category from REML variance component (VC) estimates using:

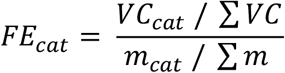

Where *VC*_*cat*_ / ∑ *VC* (proportion of VC per category, summing high and low LD bins) is equivalent to 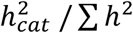 (proportion of heritability per category, summing high and low LD bins), and *m*_*cat*_ / ∑ *m* is the proportion of Variants per category. We recalculated enrichment standard errors using the Delta method for error propagation to account for uncertainty in both total and per-GRM heritability estimation:

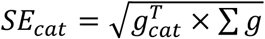

where *g* is a gradient vector describing the covariance between the per-category enrichment with each estimated VC. For the VCs combined per category (high and low LD),

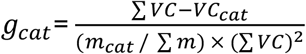

and for every other VC,

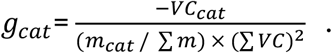

### Liability scale conversion for T2D h^2^

To estimate the heritability of binary traits, phenotypes must be implicitly converted to a continuous liability scale. Disease liability represents the likelihood for an individual to be a disease case given their liability phenotype, calculated by summing environmental and random additive genetic effects on the liability scale^66^. All individuals having liability phenotype scores exceeding a threshold T are labelled as affected.

Binary phenotypes and sample case-control ratios that do not reflect the population disease prevalence would give biased estimates because REML calculations assume that the phenotype forms a univariate Normal distribution. The observed MPH heritability estimate 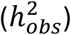 can be converted to the liability scale 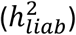 to account for the trait’s binary nature and case-control imbalances using a well-established formula (Equation 23 from Lee et al.^67^):

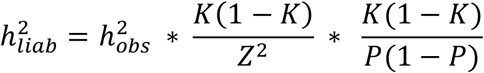

where *K*=population disease prevalence, P=sample disease prevalence, *T=*liability threshold, and *Z=*z-score at *T*. Population disease prevalences were obtained per genetic similarity group from the CDC^68^. The term 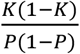 converts observed heritability onto the liability scale, and the term 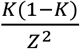 accounts for ascertainment. The converted h^2^_liab_ is used when referring to the heritability (h^2^_WGS_) of binary traits in the main text.

### Simulations

We conducted simulations in AFR to test the contribution of variant bin to heritability due to haplotype effects. Per variant bin, we randomly drew effect sizes *β* from a Gaussian distribution *β*∼*N*(0,1) scaling by MAF with effect size*(2p(1-p))^(alpha/2), alpha=-0.25. We calculated genomic scores with plink2, which we calibrated so they sum to the total observed bin h^2^_WGS_. We aggregated genomic scores for all bins and added random error corresponding to the total trait h^2^_WGS_ distributed 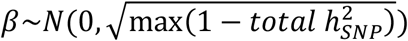 to mimic the distribution of the observed phenotype. We repeated this 10 times and META-analysed the results. We additionally simulated a condition where synonymous variants contribute 0% to heritability, and where RNA exonic variants contribute 0% to heritability.

## Notes

### Competing Interest Statement

The authors have declared no competing interest.

