## Supplementary Figures & Notes for "Quantifying heritability in functional regions of the coding and non-coding genome across diverse individuals"

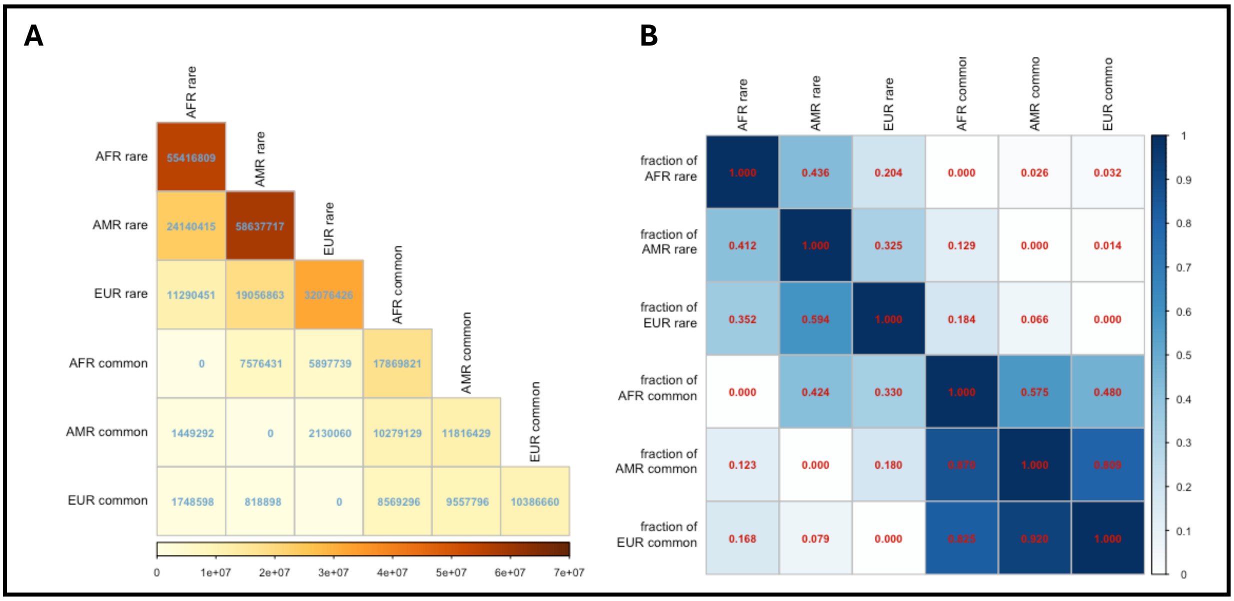


***Supplementary Figure 1: Rare (1e-4≤MAF<0.01) and common (0.01≤MAF<0.5) shared GRM variants per genetic similarity group across autosomal and X-chromosome variants.*** *Shown as* ***(A)*** *number of shared variants for rare and common-variant GRMs and* ***(B)*** *fraction of total variants per ancestry and MAF group.*

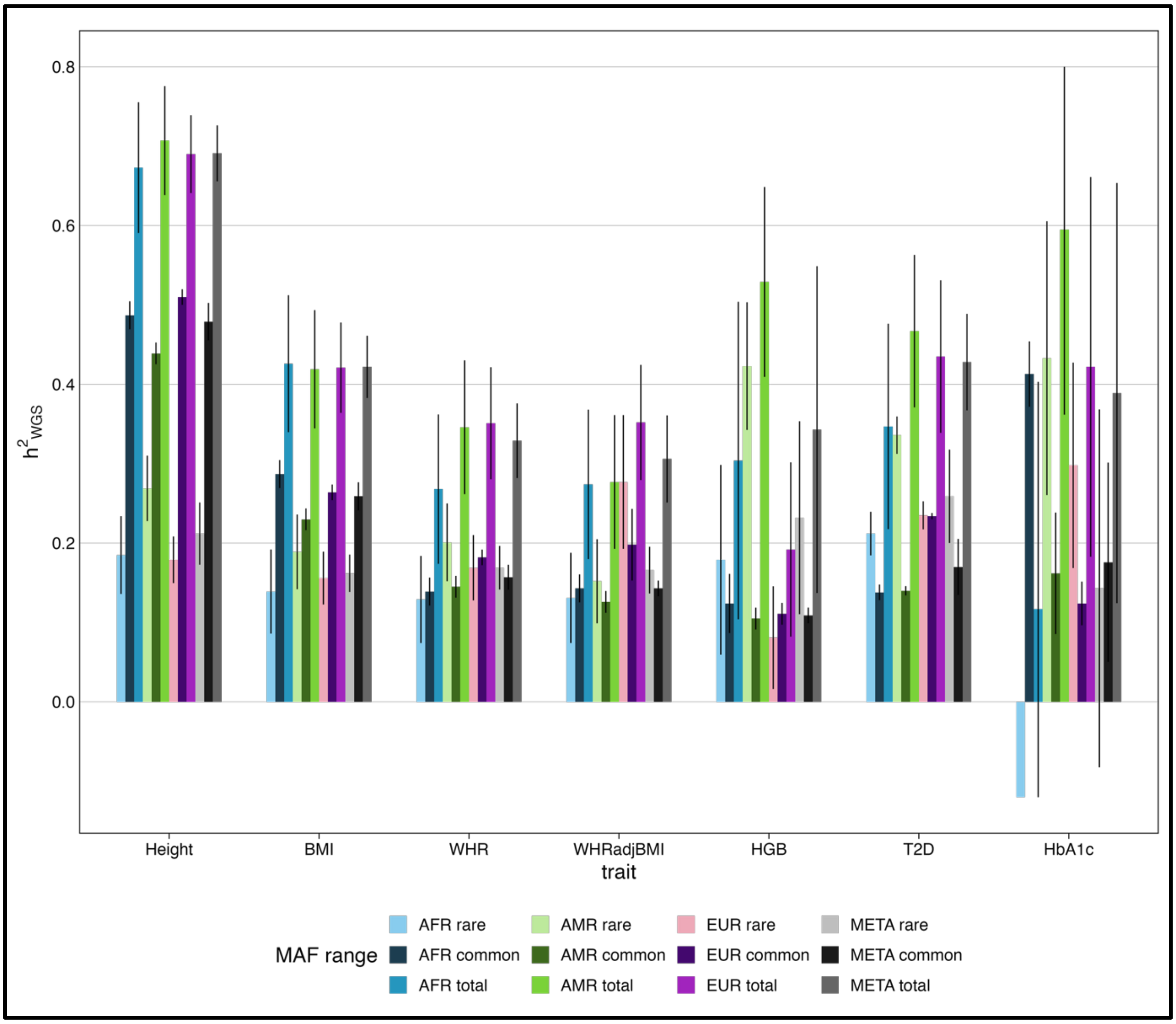


***Supplementary Figure 2: Total, rare (1e-4≤MAF<0.01), and common (0.01≤MAF<0.5) WGS heritability per trait****. Includes autosomal and chromosome X variation. Error bars represent confidence intervals (±SE*1.96) and are truncated at axis limits (-0.2≤$h^2$≤0.8).*

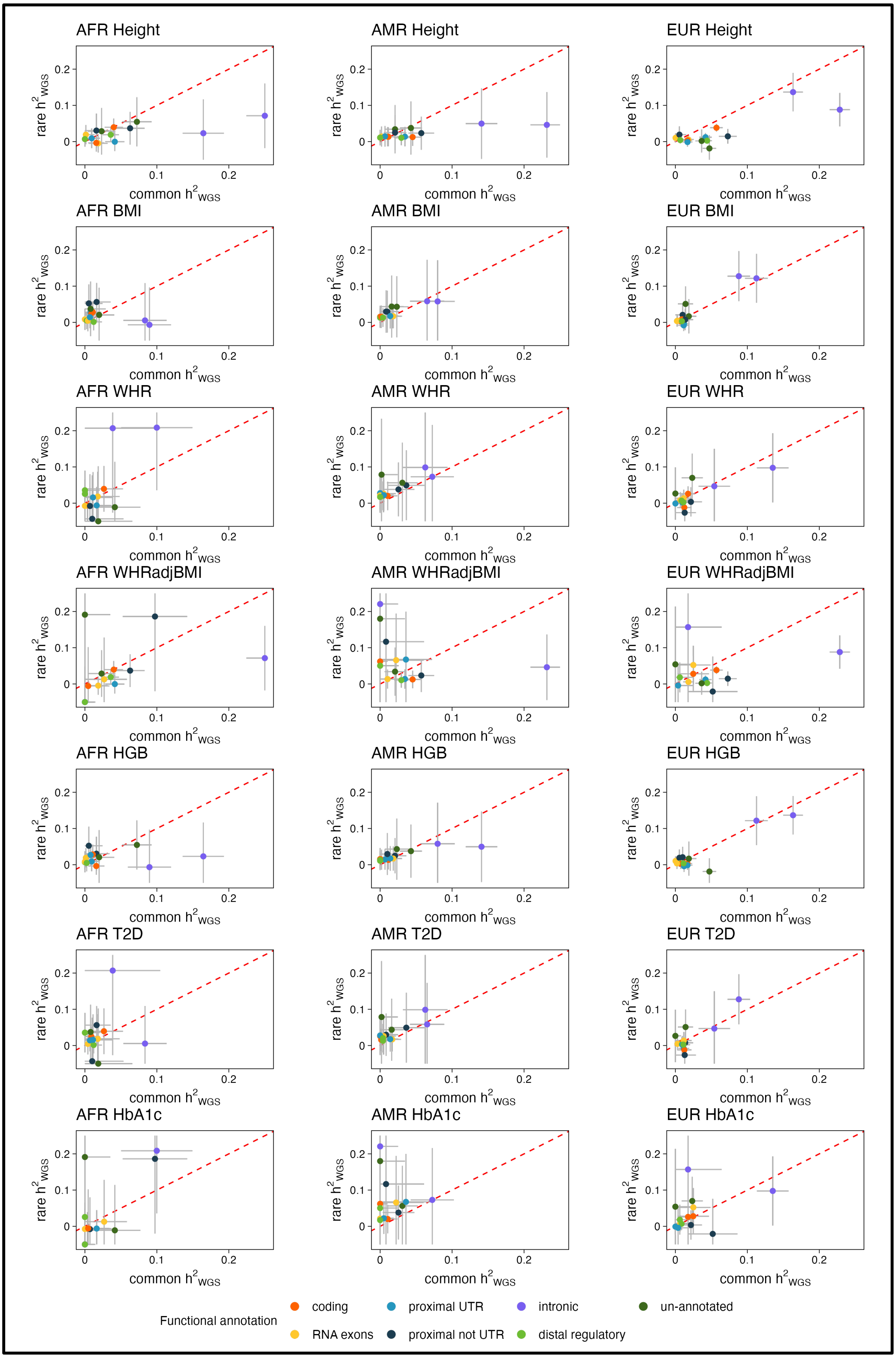


***Supplementary Figure 3: Common (0.01≤MAF<0.5) versus rare (1e-4≤MAF<0.01) $h^2_{WGS}$ per trait, annotation, and genetic similarity group.*** *Red line is y=x. Error bars represent confidence intervals (±SE*1.96) and are truncated at axis limits.*

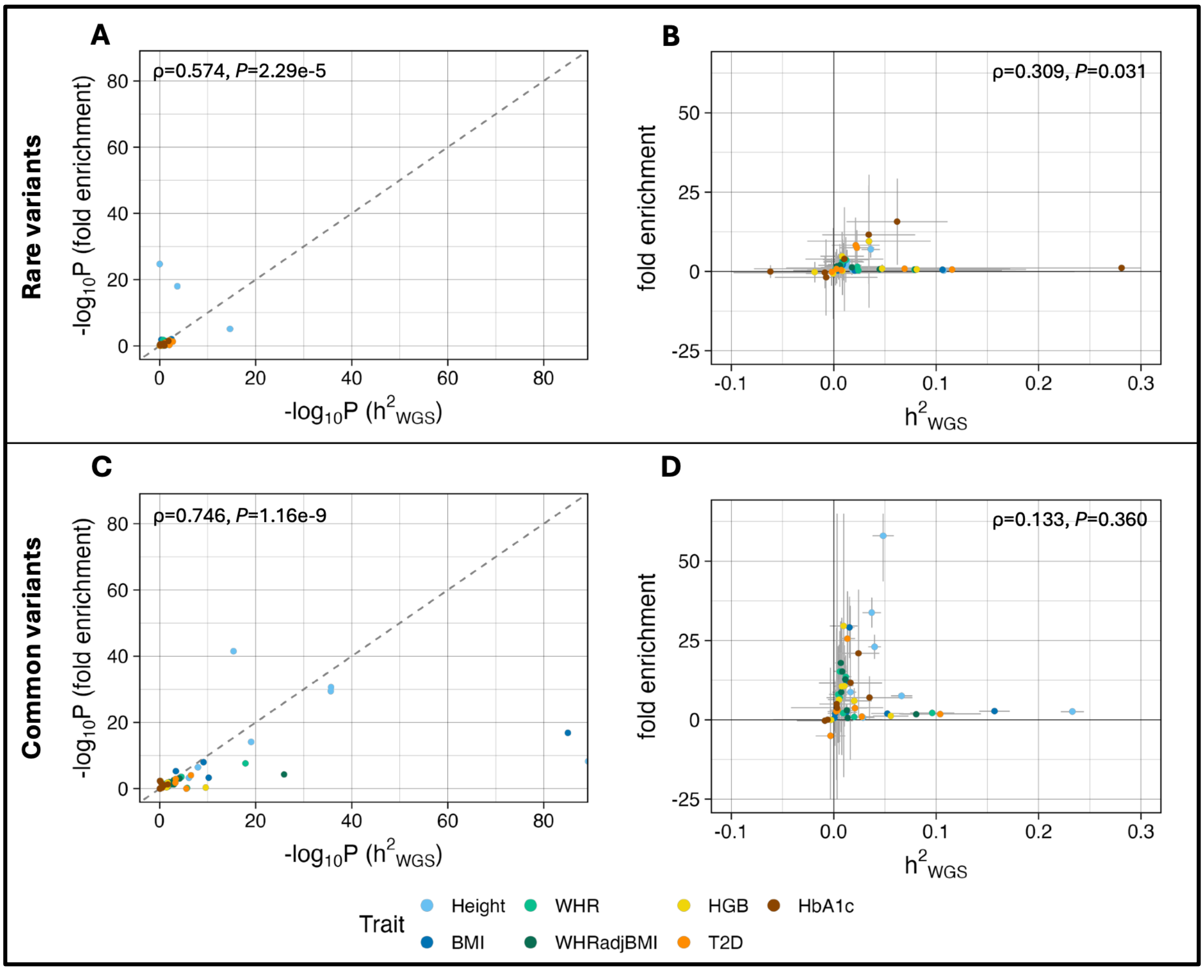


***Supplementary Figure 4: Comparing $h^2_{WGS}$ and fold enrichment per annotation category across traits for rare (A, B) and common (C, D) variants. (A, C)*** *Significance (-$\log_{10}$P) values for META-analysed $h^2_{WGS}$ and enrichment. Black diagonal line is y=x.* ***(B, D)*** *META-analysed $h^2_{WGS}$ versus fold enrichment values. Error bars represent confidence intervals (±SE*1.96) and are truncated at axis limits (-25≤FE≤65 and -0.1≤$h^2$≤0.3). Spearman correlation statistics are indicated at the top of each plot.*

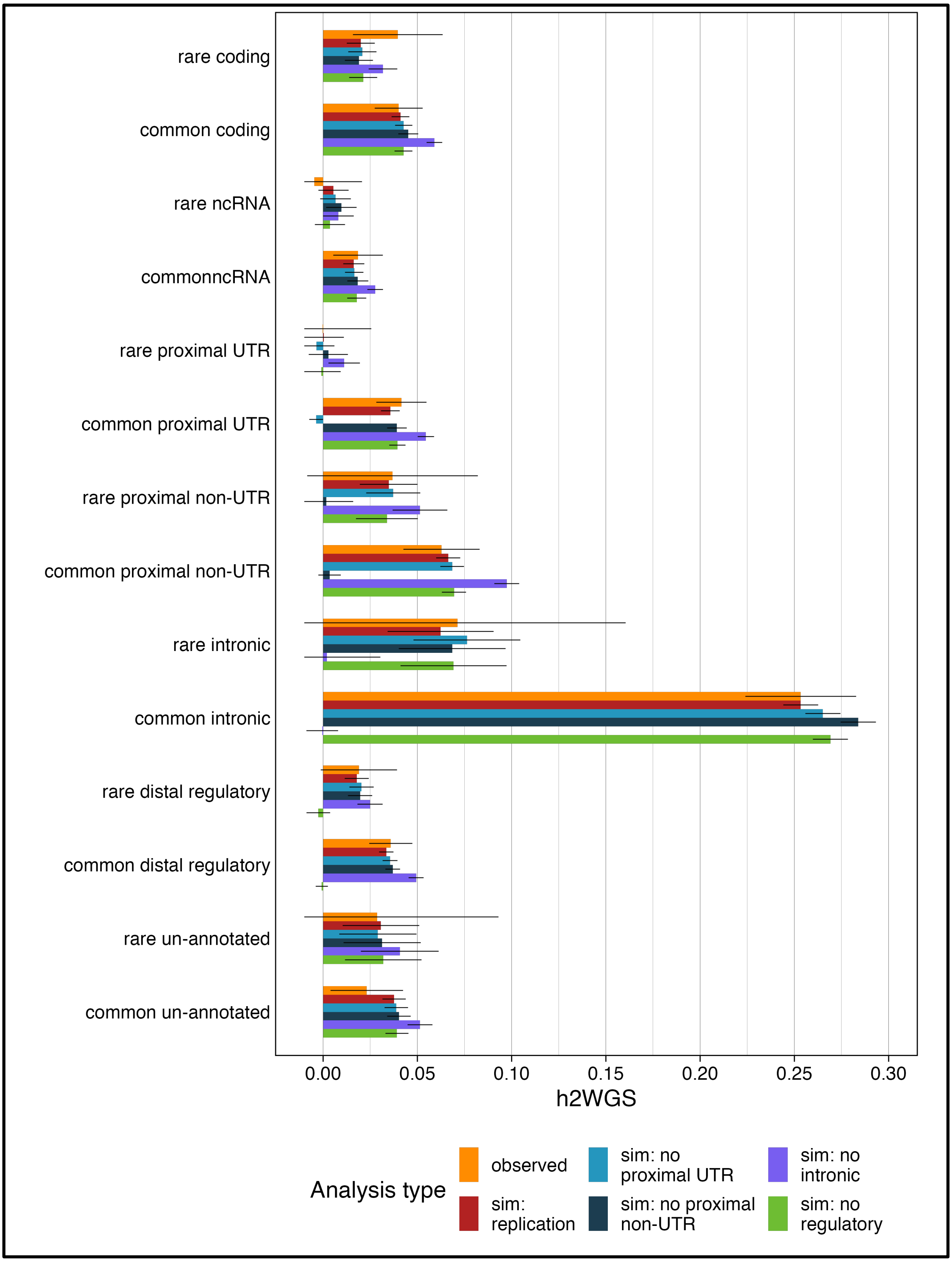


***Supplementary Figure 5: Simulated $h^2_{WGS}$ in non-coding enriched functional annotation categories****. comparing observed AFR height $h^2_{WGS}$ (orange) with simulated AFR phenotypes replicating $h^2_{WGS}$ per bin (dark red), and simulated AFR phenotypes replicating $h^2_{WGS}$ per bin while equating heritability in either proximal UTR (light blue), proximal non-UTR (dark blue), intronic (purple) or distal regulatory (green) bins to 0 while keeping total $h^2_{WGS}$=0.666. Simulation estimates per analysis type are meta-analysed across 10 repeats. Error bars represent 95% confidence intervals (±1.96*SE).*

**Supplementary Note 1: Heritability attributable to chromosome X variation**

The contribution of variants in autosomal regions versus in the X chromosome to GRMs is different, since the X-GRM is constructed under an X-specific model involving hemizygosity and sex-dependent scaling. For an autosomal variant $m$, individuals of either sex have genotypes $x_{i,m} \in \{0,1,2\}$. However, hemizygous males have different biological allele dosages $x_{i,m} \in \{0,1\}$ compared to diploid females $x_{i,m} \in \{0,1,2\}$. When building a GRM, GCTA treats male genotypes as $\in x_{i,m}\{0,2\}$ to mimic the female genotype scale[1].

Due to this dosage difference, GTCA has different variance-covariance GRM equations for female-female, male-male, and female-male pairs. For autosomes, the GRM value describing the genetic relationship between individuals *i* and *j* over an *M* number of variants *m* at allele frequency $p_m$ is:

$$GRM_{i,j} = \sum_{m=1}^{M} \frac{(x_{i,m} - 2p_m)(x_{j,m} - 2p_m)}{(2p_m(1-p_m))^{1+\alpha}} * \frac{1}{M} * \frac{1}{\sum_{m=1}^{M} 2p_m(1-p_m)}$$

With complete dosage compensation, for female-female pairs in X-chromosome the GRM value is:

$$GRM_{i,j} = \sum_{m=1}^{M} \frac{(x_{i,m} - 0.5p_m)(x_{j,m} - 0.5p_m)}{(0.5p_m(1-p_m))^{1+\alpha}} * \frac{1}{M} * \frac{1}{\sum_{m=1}^{M} 0.5p_m(1-p_m)}$$

Whereas for a male-male pair the GRM value is:

$$GRM_{i,j} = \sum_{m=1}^{M} \frac{(x_{i,m} - p_m)(x_{j,m} - p_m)}{(p_m(1-p_m))^{1+\alpha}} * \frac{1}{M} * \frac{1}{\sum_{m=1}^{M} p_m(1-p_m)}$$

And for a female-male pair, where individual *i* is female and individual *j* is male, the GRM value is:

$$GRM_{i,j} = \sum_{m=1}^{M} \frac{(x_{i,m} - 0.5p_m)(x_{j,m} - p_m)}{(1/\sqrt{2}p_m(1-p_m))^{1+\alpha}} * \frac{1}{M} * \frac{1}{\sum_{m=1}^{M} 1/\sqrt{2}p_m(1-p_m)}$$

Therefore, the underlying raw X chromosome genetic similarity between individuals is not necessarily equivalent to the X chromosome GRM value after sex-specific parametrisation, whereas these two values are equal for autosomal GRMs. In particular, variants can only be attributable to half the magnitude of genetic (and therefore phenotypic) covariance in females compared to males with complete dosage compensation.

Merging X-GRMs and autosomal GRMs would not preserve the different covariance structures of autosomal versus X-chromosome genetic relationships. Joint modelling in GREML however retains the distinct covariance structures of each variance component by estimating their contribution separately:

$$V_G = A_{auto}\sigma_{auto}^2 + A_X\sigma_X^2 + I\sigma_e^2$$

Where $V_G$ is the additive genetic variance, $A_{auto}\sigma^2_{auto}$ and $A_X\sigma^2_X$ describe the autosomal and X chromosome genetic relationship multiplied by their respective GRM covariance structure and $I\sigma^2_e$ is the environmental error term. As there are not enough variants on chromosome X to build statistically stable GRMs for functionally-partitioned heritability analyses, we only considered autosomal contributions when investigating the relative fold enrichment by genomic functional annotation.

**Supplementary Note 2: REML as an imperfect quantifier of genetic variance**

Firstly, we had reduced statistical power to detect AFR and AMR effects compared to EUR effects, especially for HGB and HbA1c which had fewer available measurements compared to the other traits, where in some cases the per-genetic similarity group sample size was N<20,000. Some stratification effects are still likely to contribute to AMR heritability estimates, even after 1) our modification of GRM calculation, 2) using MPH which is designed to accommodate cohorts with larger haplotype blocks, and 3) the inclusion of 120 genetic principal components. Previous studies have also tried adjusting for extreme population stratification by calculating rare-variant PCs to capture recent population changes[2,3] or computing local ancestry heritability which maps haplotype segments instead of individual variants to ancestral populations[4,5]. However, the inclusion of rare-variant PCs is computationally intensive with large samples and difficult without exponentially increasing estimation error[3]. Similarly, measures of local ancestry have largely been developed for and tested on the determination of archaic ancestry[22,41] or for cohorts with only two ancestral populations[5]. With our adjustments, our heritability estimates were homogenous across anthropometric traits which had larger per-ancestry sample sizes than metabolic traits.

Differences between ancestries could also be due to a number of other factors: metabolic traits are more variably measured, can change quickly during an individual's lifetime, and are more often confounded by gene-environment interactions including family environment and social determinants of health[6–8]. To investigate whether the issue is sample-size related, further heritability analysis with larger numbers of unrelated diverse individuals could more accurately and informatively detect genomic regions relative to metabolic trait heritability across non-European individuals.

Secondly, we did not correct for assortative mating in our study. Assortative mating is the preferential choice of mates based on similar phenotypic characteristics. It can lead to an increased correlation and LD between trait-increasing alleles within families and increased genetic variance in the total population, and thus artificially inflated trait heritability estimates. However, there have been conflicting theories[9,10] on whether true assortative mating (choosing mates based on phenotypic characteristics) and population stratification due to admixture (mates with correlated genotypes and phenotypes due to geographical proximity) can be accurately separated. Moreover, previous studies[11,12] which investigated the effect of assortative mating on the heritability of various traits using variant-based correlations or polygenic indexes only found evidence for moderate $h^2_{WGS}$ inflation due to assortative mating in height and educational attainment but not in other traits, including T2D, BMI, and WHR. Two studies[13,14] found evidence for assortative mating regarding T2D, but these studies only considered phenotypic and not genetic correlation between mates. For these reasons, and because our height $h^2_{WGS}$ estimates were very similar to existing literature, we did not estimate the effect of assortative mating on our results.

Thirdly, REML-based heritability alone serves as an indicator of where causal variants may lie but does not directly explain genotype-phenotype causality. Since MPH and similar methods use maximum likelihood to correlate genotypes with phenotypes and do not consider convergent gene pathways and pleiotropic effects, they are not a direct indication of genotype-phenotype links even in some cases when these specific effects are captured in separate variance components (GRMs). For example, our results attribute a large part of protein-coding height and BMI heritability to common synonymous variants, for which there is little GWAS-based association evidence. Accounting for LD-related error through GREML-LDMS or simulations does not completely remove this observation, so it is unclear whether the result captures true aggregated small synonymous variant effects or whether the result is biased in favour of common synonymous bins as they are more stable statistical predictors for trait heritability than the other fewer, lower MAF, more poorly genotyped coding regions. This result, combined with the fact that polygenic phenotypes display several levels of biological organisation via pleiotropic metabolic pathways and tissue-specific expression differences, warrants further investigation via burden testing, functional mapping, and tissue-specific or single-cell RNAseq investigations.
